# Making Plants Smell Like Moths: *Nicotiana benthamiana* release moth pheromone alcohol, aldehyde and acetate upon transient expression of biosynthetic genes of different origin

**DOI:** 10.1101/2021.09.10.459774

**Authors:** Yi-Han Xia, Bao-Jian Ding, Shuang-Lin Dong, Hong-Lei Wang, Per Hofvander, Christer Löfstedt

**Affiliations:** Department of Biology, Lund University, Sölvegatan 37, SE-22362, Lund, Sweden; Department of Plant Breeding, Swedish University of Agricultural Sciences, P.O. Box 101, SE-23053 Alnarp, Sweden; Education Ministry Key Laboratory of Integrated Management of Crop Diseases and Pests, College of Plant Protection, Nanjing Agricultural University, CN-210095, Nanjing, China; Department of Biology and Biological Engineering, Chalmers University of Technology, Kemivägen 4, SE-41296, Gothenburg, Sweden

**Keywords:** Functional characterization, fatty acyl desaturases, fatty acyl reductase, alcohol oxidation, acetyltransferase, heterologous expression systems, pheromone-releasing plants

## Abstract

Using genetically modified plants as natural dispensers of insect pheromones may eventually become part of a novel strategy for integrated pest management. In the present study, we first characterized essential functional genes for sex pheromone biosynthesis in the rice stem borer *Chilo suppressalis* (Walker) by heterologous expression in *Saccharomyces cerevisiae* and *Nicotiana benthamiana*, including two desaturase genes *CsupYPAQ* and *CsupKPSE*, and a reductase gene *CsupFAR2*. Subsequently, we co-expressed *CsupYPAQ* and *CsupFAR2* together with the previously characterized moth desaturase *AtrΔ11* in *N. benthamiana*. This resulted in the production of (*Z*)-11-hexadecenol together with (*Z*)-11-hexadecenal, the major pheromone component of *C. suppressalis*. Both compounds were collected from the transformed *N. benthamiana* headspace volatiles using solid phase microextraction. We finally added the expression of a yeast acetyltransferase gene *ATF1* and could then confirm also (*Z*)-11-hexadecenyl acetate release from the plant. Our results pave the way for stable transformation of plants to be used as biological pheromone sources in different pest control strategies.

## Introduction

Moths rely strongly on sex pheromones for mate communication. Synthetic pheromones have been used for monitoring, mass trapping and mating disruption in Integrated Pest Management (IPM) for several decades (Trematerra 1997; Witzgall et al. 2010) as environmentally friendly alternatives or complements to conventional insecticides. They are species-specific and non-toxic, and the risk of pests evolving resistance to their own pheromone is very low. Compared to current standard approaches to pheromone synthesis (Mori 2010), the use of biological factories for pheromone productions may have several advantages, allowing cost-efficient production of moderate to large quantities of pheromones with high purity and a minimum of waste. A series of proof-of-concept studies have clearly demonstrated the potential of producing moth pheromones in both plant and yeast factories (Hagström et al. 2013b; Ding et al. 2014; Xia et al. 2020; Holkenbrink et al. 2020) to replace conventionally produced pheromones in existing systems for pheromone-based pest control. However, under certain circumstances it may be advantageous to cultivate plants that actually release the pheromone volatiles in the field rather than accumulates them for harvest. Such pheromone-releasing plants could hypothetically and depending on context either protect themselves against moth pests by causing mating disruption or be cultivated together with other harvest crops for the purpose of attraction or mating disruption.

Deciphering the molecular mechanism of pheromone biosynthesis in moth species can provide a functional gene pool for producing customized pheromones in heterologous systems. Approximately 75% of the identified moth pheromones belong to the so-called Type I sex pheromones; C_10_-C_18_ alcohols, acetates, or aldehydes that are biosynthesized from palmitic and stearic acid by consecutive steps of fatty acyl desaturation, chain-shortening or chain-elongation, reduction, and final modification (Löfstedt et al. 2016). Specific enzymes are used for each catalytic step. Fatty acyl desaturases (FADs) are the first essential enzymes that introduce double bonds in specific positions of carbon chains with strict regioselectivity and stereoselectivity. Fatty acyl reductases (FARs), which are responsible for reducing fatty acyl to fatty alcohols, are essential for forming functional groups of the fatty acid derivatives (Löfstedt et al. 2016). The genes encoding the essential enzymes involved in pheromone fatty acyl desaturation and reduction have been functionally characterized in many moth species (Ding et al. 2011; Hao et al. 2002a; 2002b; Knipple et al. 1998, 2002; Liu et al. 2002, 2004; Rodríguez et al. 2004; Roelofs et al. 2002; Rosenfield et al. 2001; Serra et al. 2007; Wang et al. 2010; Xia et al. 2019; Lassance et al. 2010; Moto et al. 2003; Antony et al. 2016). Characterization of FAD and FAR genes with different substrate and product specificities remains important for the production of tailored moth pheromones in heterologous systems.

The fatty alcohols may be actual pheromone components, or they may be further functionalized by the conversation of fatty alcohols into esters (Clinkenbeard et al. 1973) or aldehydes (Teal and Tumlinson 1986; Fang et al. 1995; Hoskovec et al. 2002). Although *in vivo* labeling experiments have confirmed these biological reactions, no enzymes catalyzing the reactions in the pheromone glands have been identified and cloned from any insect species. In order to produce acetate pheromones in plants, Ding et al. (2014) explored the possibility of using an acetyltransferase gene *EaDAcT* cloned from burning bush, *Euonymus alatus*. Transient expression resulted in the production of (*Z*)-11-hexadecenyl acetate (Z11-16:OAc), but the efficiency was low. Subsequently, Ding et al. (2016) characterized a yeast acetyltransferase gene *ATF1* which efficiently acetylates insect pheromone alcohols into acetates in yeast. However, the activity of *ATF1* in plants remains to be explored.

The rice stem borer, *Chilo suppressalis* (Walker) (Lepidoptera: Crambidae), boring the stems of their host plants, is an infamous rice pest in East Asia, India and Indonesia, causing great production reduction in rice crops (Zhu et al. 2007). In the 1970s and 1980s, the sex pheromone of female *C. suppressalis* was identified as a mixture of (*Z*)-11-hexadecenal (Z11-16:Ald), (*Z*)-13-octadecenal (Z13-18:Ald) (Nesbitt 1975; Ohta et al. 1976) and (*Z*)-9-hexadecenal (Z9-16:Ald) at the ratio of 100:13:11 (Tatsuki 1983). In the present study, we functionally characterized several candidate genes likely to be involved in pheromone biosynthesis in *C. suppressalis* by heterologous expression in yeast and plant platforms, using the transcriptome data reported by Xia et al. (2015). We produced *N. benthamiana* genetically modified for expression of the functionally characterized *C. suppressalis* Δ11 desaturase *CsupYPAQ*, the fatty acyl reductase *CsupFAR2* and the yeast acetyltransferase *ATF1* that released a mixture of (*Z*)-11-hexadecenol (Z11-16:OH), Z11-16:OAc and Z11-16:Ald. Our study contributes additional genes to the pool of key enzymes available for biotechnological production of moth pheromones, and more importantly it is a significant step forward in construction of genetically modified plants to be used as natural dispensers of insect pheromones as part of IPM strategies for pest control.

## Results

### *CsupYPAQ* and *CsupKPSE* are the functional FAD genes involved in pheromone biosynthesis in *Chilo suppressalis*

The *C. suppressalis* FAD-like genes *CsupYPAQ* and *CsupKPSE* displayed 1038 nt and 1059 nt ORFs that translated into 346 and 353 aa-proteins, respectively. In a phylogenetic analysis of lepidopteran FADs, *CsupYPAQ* clustered into the Δ11/Δ10/multifunctional FAD subfamily, whereas *CsupKPSE* fell into the Δ9 (C_16_>C_18_) FAD clade (Fig. 1).

**Figure 1.**
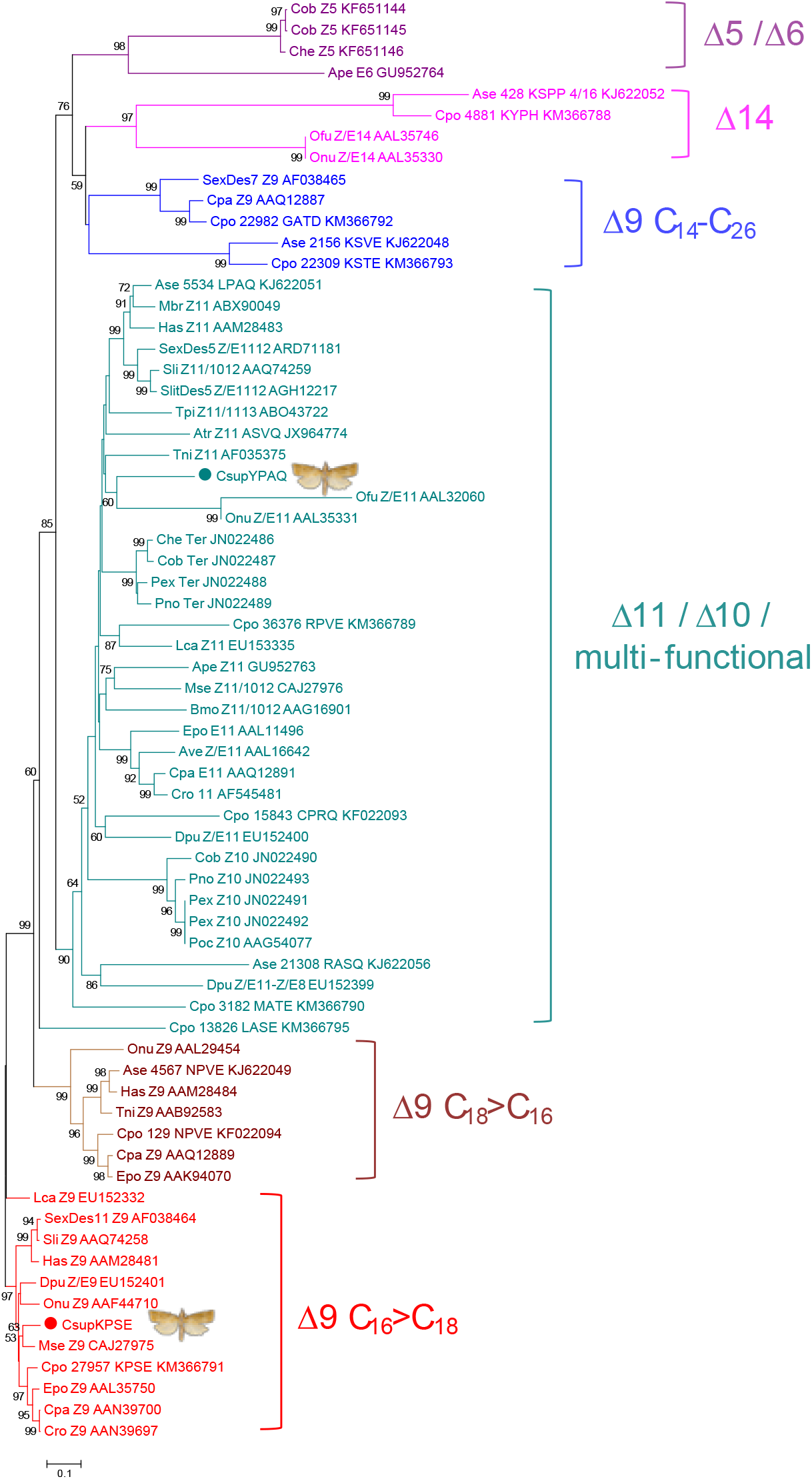
Phylogenetic tree of fatty acyl desaturases (FADs). The FAD tree was constructed by using amino acid sequences of lepidopteran FADs. The FADs functionally characterized in previous studies are classified into six subfamilies: Δ9 FAD subfamilies usually involved in normal fatty acid metabolism with preference for C_16_ (C_16_>C_18_) or C_18_ (C_18_>C_16_); the Δ11/Δ10/multifunctional FAD subfamily; the Δ9 FAD subfamily with preference for C_14_-C_26_, involved in pheromone biosynthesis; and the Δ14 and Δ5/Δ6 FAD subfamilies that introduce double bonds into unusual positions in saturated fatty acids. The predicted FAD genes *CsupYPAQ* and *CsupKPSE* from *C. suppressalis* are marked by blue and red filled circles, respectively.

The functional expression of *CsupYPAQ* and *CsupKPSE* indicated their involvement in pheromone biosynthesis. GC/MS analysis of yeast fatty acids showed that the yeast expressing *CsupYPAQ* produced a high amount of (*Z*)-11-hexadecenoic acid (Z11-16:acid) (Fig. 2a), while yeast expressing *CsupKPSE* produced high amounts of oleic acid (Z9-18:acid) and (*Z*)-9-hexadecenoic acid (Z9-16:acid) (Fig. 2b). Compared to the yeast expression results, *CsupYPAQ* showed a similar function in *N. benthamiana* (Fig. 3a). However, *CsupKPSE* expressed in *N. benthamiana* did not produce observable extra Z9-18:acid but only a very high amount of Z9-16:acid (Fig. 3b). In the wild type *N. benthamiana*, none of the monounsaturated potential pheromone precursors was produced (Fig. 3c).

**Figure 2.**
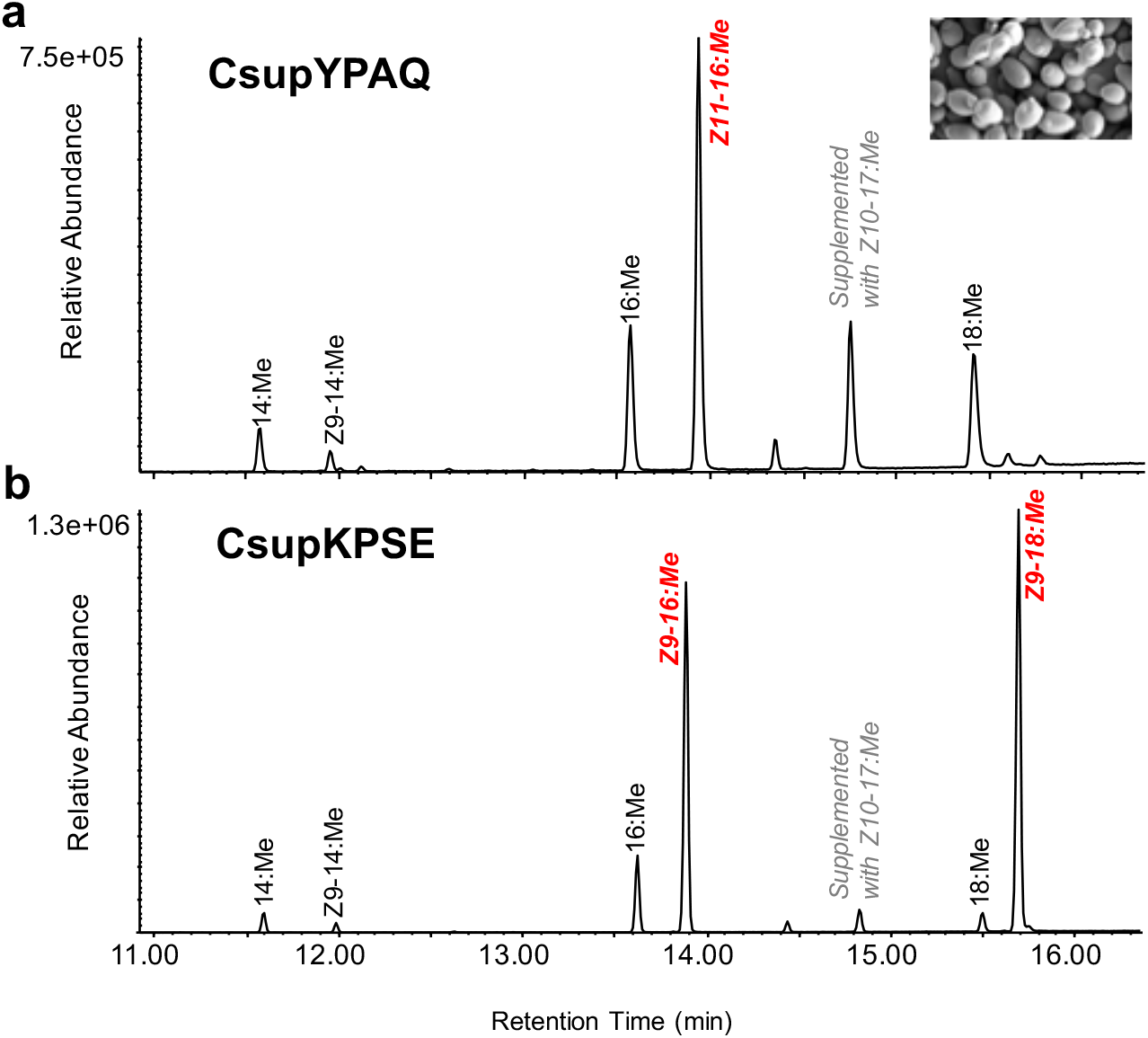
Heterologous expression of fatty acyl desaturase candidates from *Chilo suppressalis* in Δ*ole1*/*elo1 Saccharomyces cerevisiae*. GC/MS analysis of fatty acid methyl ester profiles of yeast expressing the *CsupYPAQ* or *CsupKPSE*. The compounds produced from the introduced desaturases are labelled in red italics. Methyl (*Z*)-10-heptadecanoate (Z10–17:Me) was supplemented as nutrition to all the incubations.

**Figure 3.**
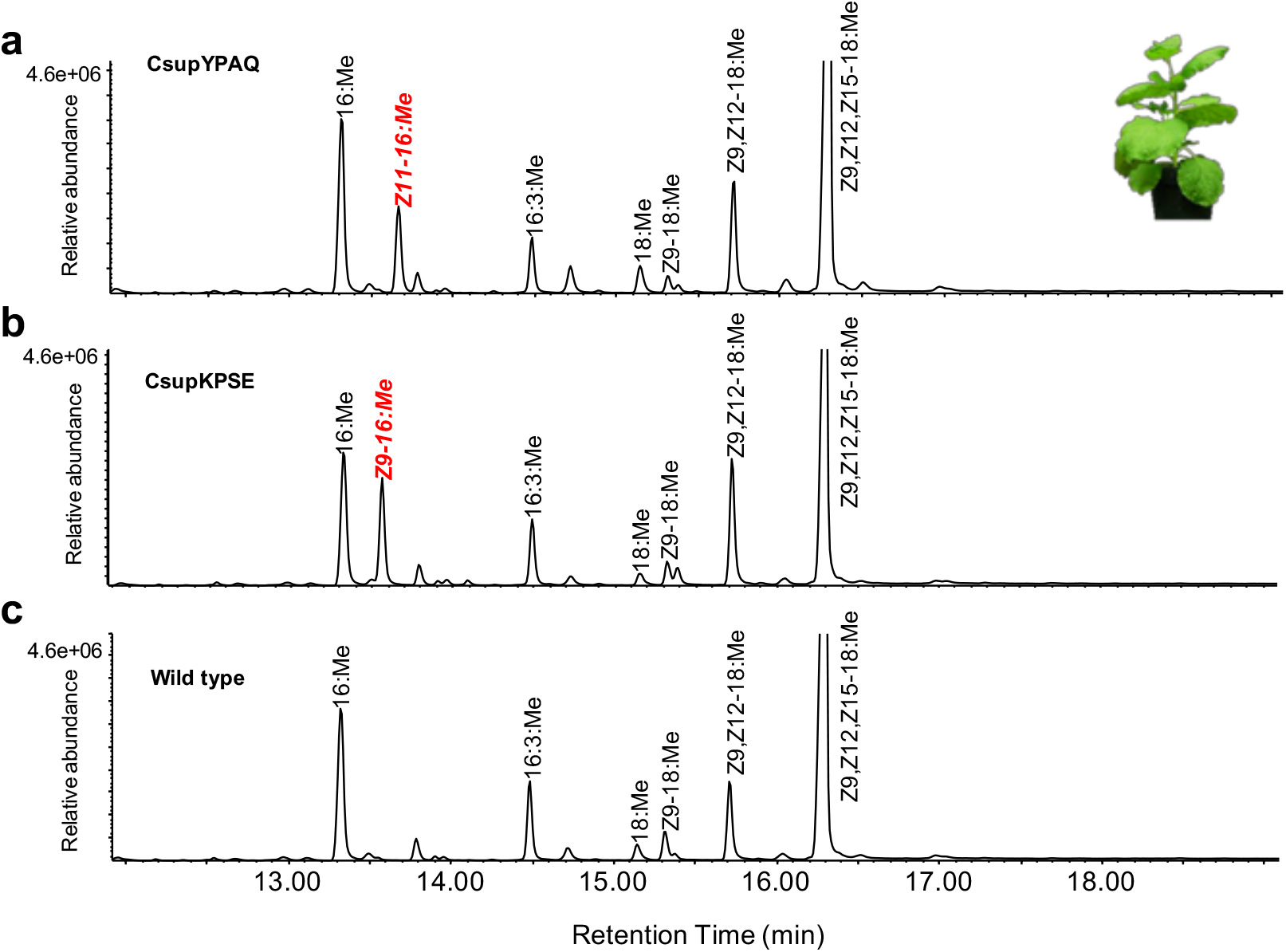
Heterologous expression of fatty acyl desaturase candidates from *Chilo suppressalis* in *Nicotiana benthamiana*. GC/MS analysis of fatty acid methyl ester profiles of a) plant leaves expressing *CsupYPAQ*, and b) plant leaves expressing *CsupKPSE*, and c) wild type leaves. Native compounds from plant are labelled in black and the compounds produced from the introduced desaturases are labelled in red italics.

### *CsupFAR2* is the FAR involved in pheromone biosynthesis in *Chilo suppressalis*

The gene candidate *CsupFAR2* encompassed an ORF of 1404 nt, which corresponded to a 468 aa-protein. The phylogenetic analysis showed that *CsupFAR2* could be classified as a pgFAR (Fig. 4).

**Figure 4.**
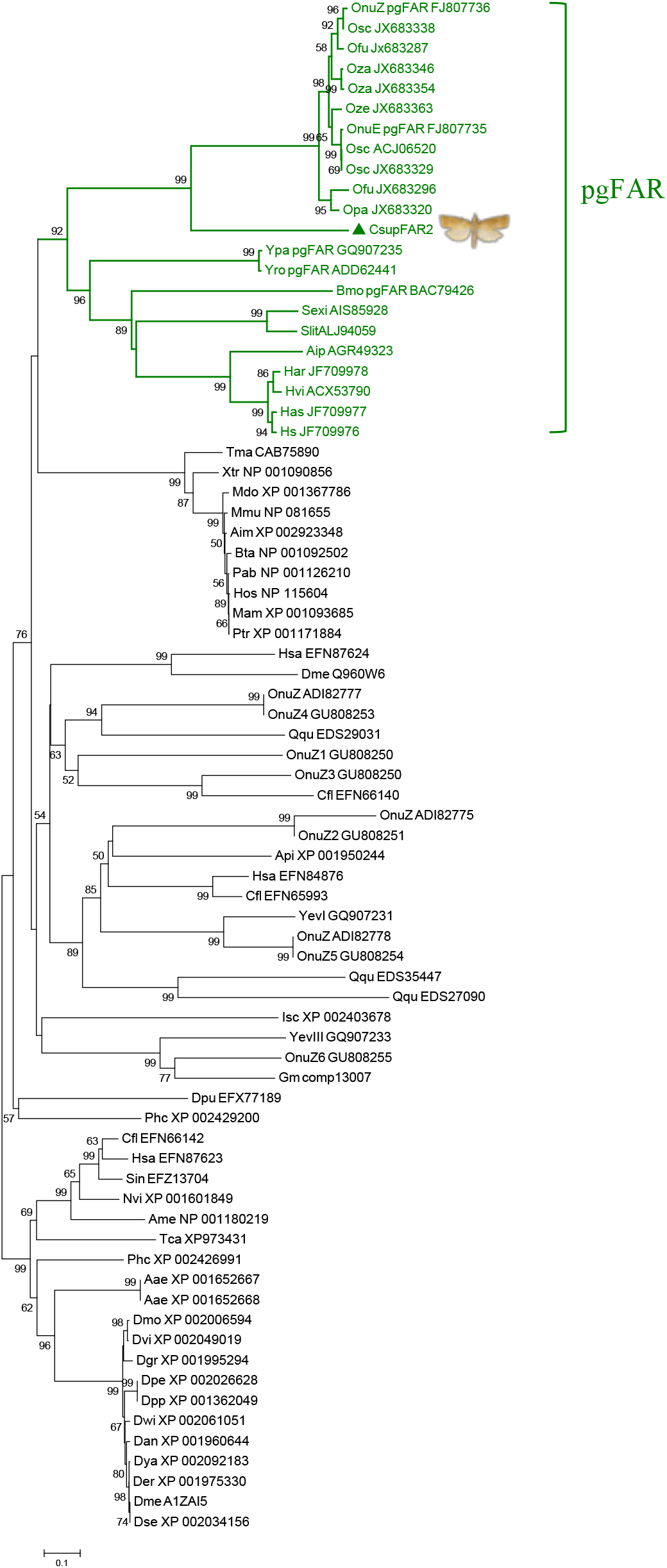
Phylogenetic tree of fatty acyl reductases (FARs). The tree is constructed from mammal, arthropod and Lepidoptera FARs using amino acid sequences. The pgFAR clade, which contains previously functionally characterized FARs involved in moth pheromone biosynthesis, is shown in green and marked by a bracket. The predicted *C. suppressalis* fatty acyl reductase *CsupFAR2* is marked by a triangle.

A co-expression construct of *CsupYPAQ*-*CsupFAR2* was produced (ca. 3500 nt) with the *Gal1* promoter upstream of *CsupFAR2*. The functional assays of *CsupFAR2* demonstrated that it reduced the *C. suppressalis* pheromone precursors Z11-16:acid, Z9-16:acid and Z13-18:acid to the corresponding fatty alcohols (Fig. 5-6).

**Figure 5.**
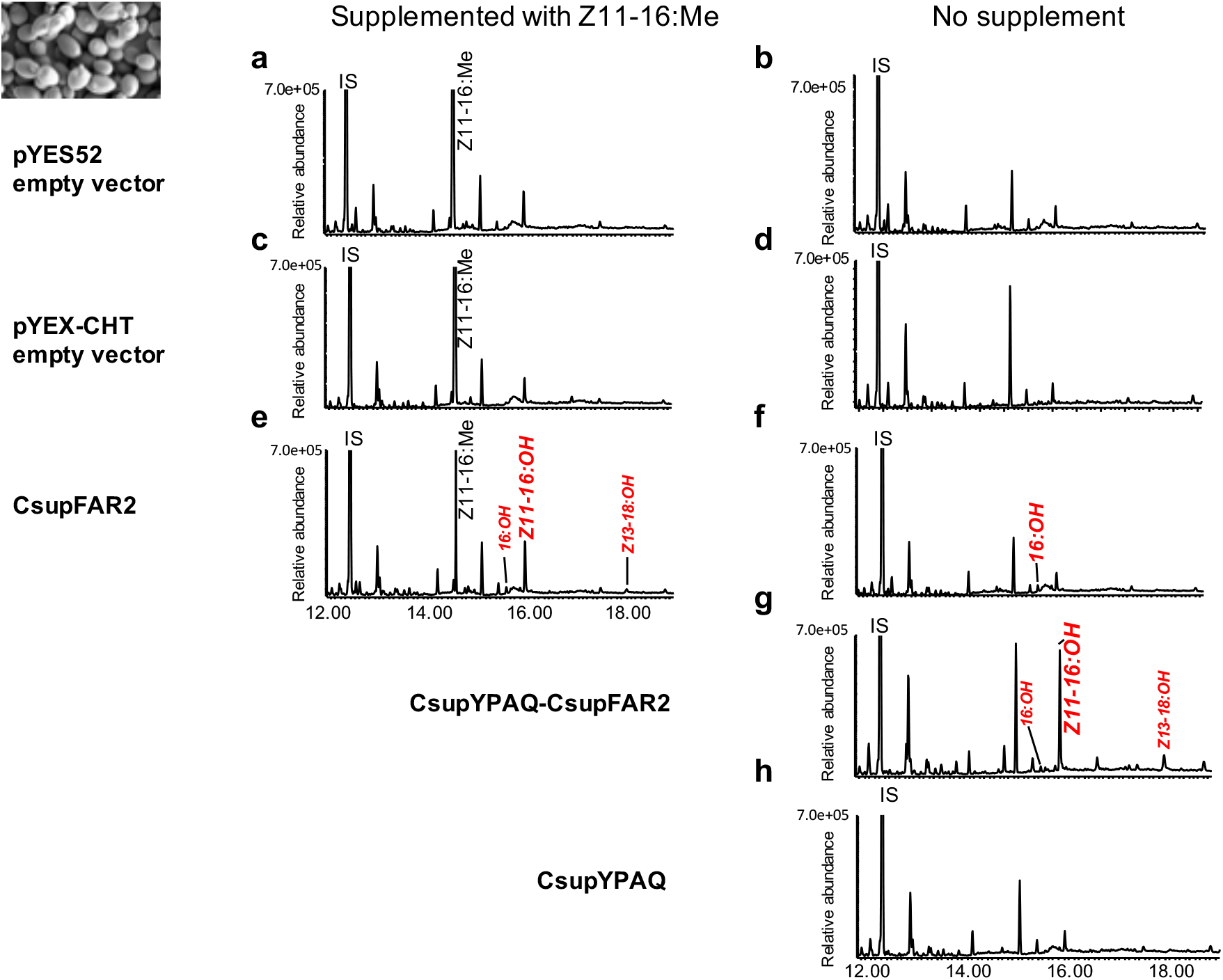
Heterologous expression of the fatty acyl reductase candidate gene *CsupFAR2* and the desaturase gene *CsupYPAQ* in *Saccharomyces cerevisiae INVSc* strain. *CsupFAR2* is constructed in the galactose inducible vector pYES52 and *CsupYPAQ*-*CsupFAR2* is constructed in the Cu^2+^ inducible vector pYEX-CHT. GC/MS analysis of fatty alcohol profiles of yeast harboring the a) pYEX52 empty vector supplemented with Z11-16:Me; b) pYES52 empty vector without supplement; c) pYEX-CHT empty vector supplemented with Z11-16:Me; d) pYEX-CHT empty vector without supplement; e) *CsupFAR2* in pYES52 supplemented with Z11-16:Me; f) *CsupFAR2* in pYES52 without supplement; g) *CsupYPAQ-CsupFAR2* and h) *CsupYPAQ* in pYEX-CHT without supplement. Fatty alcohols produced from the introduced genes are shown in red and italics.

The yeast with the transformed empty vectors of Gateway adjusted pYES52 (Fig. 5a-b) and pYEX-CHT (Fig. 5c-d) did not produce fatty alcohol, while the yeast expressing *CsupFAR2* or *CsupYPAQ-CsupFARs* produced Z11-16:OH, (*Z*)-13-octadecenol (Z13-18:OH), and palmityl alcohol (16:OH) (Fig. 5e-g). With the latter construct (*CsupYPAQ*-*CsupFAR2*), yeast produced a higher amount of target compounds (Fig. 5g). In the negative control, the yeast expressing *CsupYPAQ* did not produce fatty alcohols (Fig. 5h). Besides Z11-16:OH, Z13-18:OH and 16:OH, no other fatty alcohol species were detected in the yeast (Fig. 5).

In the *N. benthamiana* expression system, *CsupFAR2* was demonstrated to be active with a range of substrates including saturated, monounsaturated and polyunsaturated fatty acids. The leaves expressing *CsupFAR2* reduced a high amount of 16:acid and a minor amount of α-linolenic acid (Z9,Z12,Z15-18:acid) to the corresponding 16:OH and linolenyl alcohol (Z9,Z12,Z15-18:OH) (Fig. 6a).

**Figure 6.**
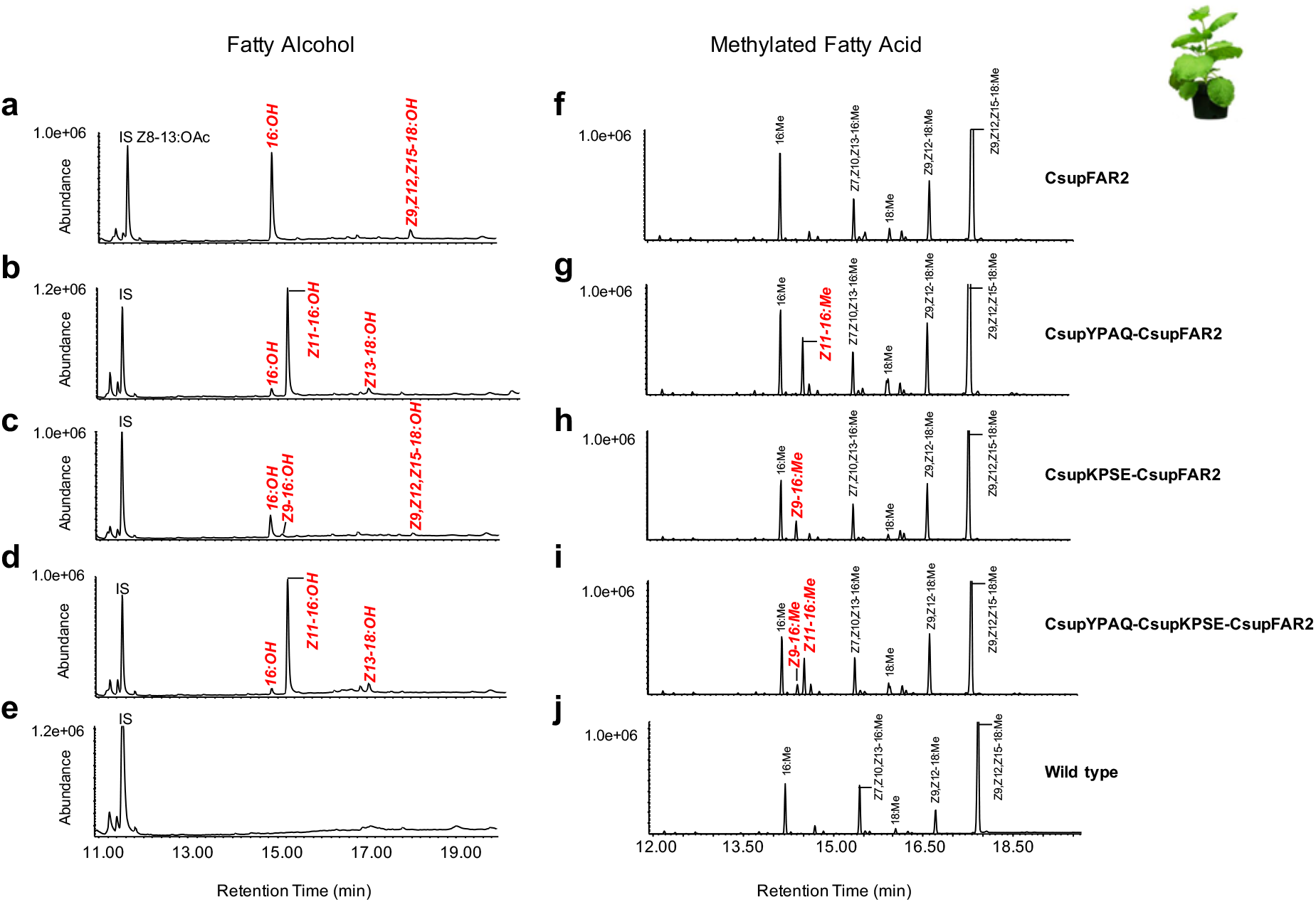
Heterologous expression of fatty acyl reductase candidate gene *CsupFAR2* and fatty acyl desaturase genes *CsupYPAQ* and *CsupKPSE* in *Nicotiana benthamiana*. GC/MS analysis of fatty alcohol (a-e) and corresponding fatty acid (f-g) profiles of plant leaves expressing the a) and f) *CsupFAR2*; b) and g) *CsupYPAQ-CsupFAR2*; c) and h) *CsupKPSE-CsupFAR2*; d) and i) *CsupYPAQ-CsupKPSE-CsupFAR2*, and wild type (e and g). Fatty alcohols produced by the introduced genes are shown in red. 18 µg of Z8-13:OAc per gram fresh leaf was added during the extraction as internal standard.

However, it did not reduce other plant-derived fatty acids such as linoleic acid (Z9,Z12-18:acid), palmitolinolenic (7,10,13-16:acid), Z9-18:acid or other saturated fatty acids than 16:acid in detectable amounts. Furthermore, the leaves produced plenty of Z11-16:OH when *CsupYPAQ* was co-introduced with *CsupFAR2*. This gene combination resulted in a ten-fold higher amount of Z11-16:OH than 16:OH and Z13-18:OH (Fig. 6b). When *CsupKPSE*-*CsupFAR2* was co-expressed in the plant, in addition to 16:OH and linolenyl alcohol, a minor amount of (*Z*)-9-hexadecenol (Z9-16:OH) was produced (Fig. 6c). The plant co-expressing multiple genes *CsupYPAQ*-*CsupKPSE*-*CsupFAR2* showed similar alcohol production compared to the plant expressing *CsupYPAQ*-*CsupFAR2* (Fig. 6b and 6d), but Z9-16:acid was only produced in the plants expressing the *CsupKPSE* (Fig. 6g and 6i). The wild type plant did not produce any of the alcohols mentioned above (Fig. 6e).

### No functional alcohol oxidase genes characterized from *Chilo suppressalis*

A homology search yielded five fatty alcohol oxidase/dehydrogenase transcript candidates (*CsupFAO_15570, CsupFAO_9572, CsupADH_10975, CsupADH_14583, CsupADH_17286*) from *C. suppressalis* that were tested in this study (Table 1). The fatty alcohol oxidase (FAO) gene candidates encompassed ORFs around 1900 nt and had the highest amino acid identity of 26.5% between *CsupFAO_15570* and an *FAO* (XM_500864) from the yeast *Yarrowia lipolytica*. The alcohol dehydrogenase (ADH) gene candidates encompassed ORFs around 1100 nt and had the highest amino acid identity of 68.6% between *CsupADH_14583* and an *ADH* (NP_741507) from the nematode *Caenorhabditis elegans*.

**Table 1.** Fatty alcohol oxidase/dehydrogenase candidates from *Chilo suppressalis* tested in this study.

| Gene ID | Csup_PG RPKM | ORF (bp) | Source of references |  |  |  |
| --- | --- | --- | --- | --- | --- | --- |
|  |  |  | Name | Acc. number | Species | Identity (%) |
| **CsupFAO_15570** | 23.6 | 1896 | Alcohol oxidase | XM_500864 | [*Yarrowia lipolytica*] | 26.5 |
|  |  |  | Alcohol oxidase | BAO74119 | [*Starmerella bombicola*] | 13.6 |
| **CsupFAO_9572** | 28.3 | 1893 | Alcohol oxidase | XM_500864 | [*Yarrowia lipolytica*] | 26.1 |
|  |  |  | Alcohol oxidase | BAO74119 | [*Starmerella bombicola*] | 14.3 |
| **CsupADH_10975** | 8.8 | 1077 | Alcohol dehydrogenase | NP_000660 | [*Homo sapiens*] | 26.3 |
|  |  |  | Formaldehyde dehydrogenase | NP_524310 | [*Drosophila melanogaster*] | 34.9 |
|  |  |  | Alcohol dehydrogenase | NP_001075997 | [*Equus caballus*] | 25.1 |
|  |  |  | Bifunctional Alcohol dehydrogenase S288C | NP_010113 | [*Saccharomyces cerevisiae*] | 24.3 |
|  |  |  | Alcohol dehydrogenase | NP_031435 | [*Mus musculus*] | 24.0 |
|  |  |  | Alcohol dehydrogenase | NP_177837 | [*Arabidopsis thaliana*] | 30.9 |
|  |  |  | Alcohol dehydrogenase class3 | NP_571924 | [*Danio rerio*] | 40.2 |
|  |  |  | Alcohol dehydrogenase class3 | NP_741507 | [*Caenorhabditis elegans*] | 37.7 |
|  |  |  | Formaldehyde dehydrogenase | XP_001693934 | [*Chlamydomonas reinhardtii*] | 34.5 |
| **CsupADH_14583** | 28.1 | 1131 | Alcohol dehydrogenase | NP_000660 | [*Homo sapiens*] | 55.3 |
|  |  |  | Formaldehyde dehydrogenase | NP_524310 | [*Drosophila melanogaster*] | 72.1 |
|  |  |  | Alcohol dehydrogenase | NP_001075997 | [*Equus caballus*] | 55.3 |
|  |  |  | Bifunctional Alcohol dehydrogenase S288C | NP_010113 | [*Saccharomyces cerevisiae*] | 60.2 |
|  |  |  | Alcohol dehydrogenase | NP_031435 | [*Mus musculus*] | 56.3 |
|  |  |  | Alcohol dehydrogenase | NP_177837 | [*Arabidopsis thaliana*] | 51.7 |
|  |  |  | Alcohol dehydrogenase class3 | NP_571924 | [*Danio rerio*] | 77.7 |
|  |  |  | Alcohol dehydrogenase class3 | NP_741507 | [*Caenorhabditis elegans*] | 68.6 |
|  |  |  | Formaldehyde dehydrogenase | XP_001693934 | [*Chlamydomonas reinhardtii*] | 62.3 |
| **CsupADH_17286** | 284 | 1134 | Formaldehyde dehydrogenases | XP_001693934 | [*Chlamydomonas reinhardtii*] | 21.9 |
| **HzeaADH7** | 241.5 | 1260 | - | - | - | - |

We then individually co-expressed the five FAO and ADH-like genes and an additional ADH gene from *Helicoverpa zea HzeaADH7*, which was reported as an ADH-highly like gene with PG abundant expression in pheromone glands (Dou et al. 2019), in *N. benthamiana* leaves together with *CsupYPAQ* and *CsupFAR2* in various combinations (Table 2).

**Table 2.**
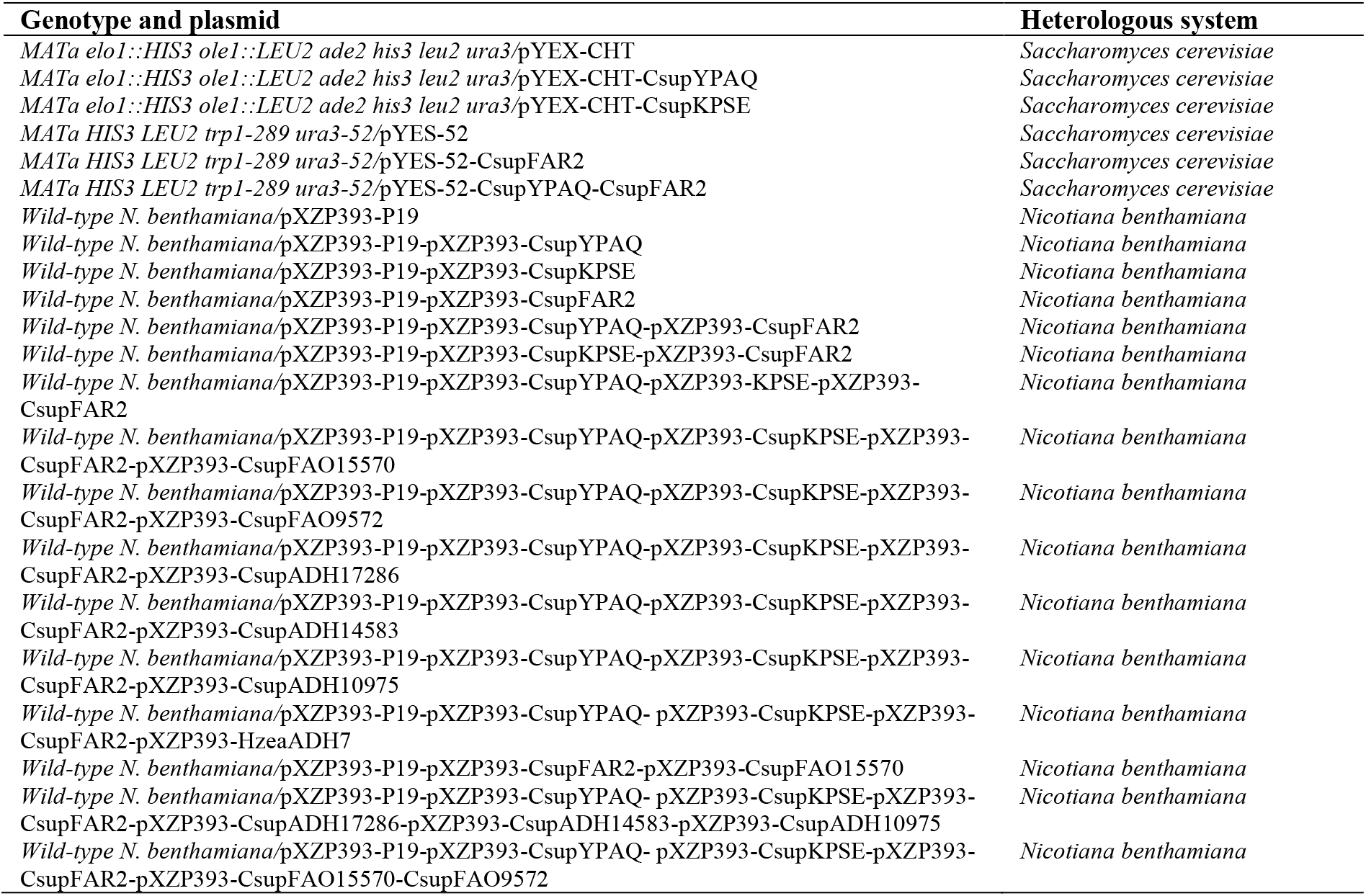
Expression vectors used for functional assays in this study.

However, no results indicating the expected activity were obtained with any of the FAO/ADH gene candidates. In the GC/MS analysis of the leaf extracts, only a small peak of Z11-16:Ald was detected in the leaf samples when *CsupYPAQ, CsupKPSE, CsupFAR2* and *Csup15570* were co-expressed in the plant (Fig. 7a) and the control leaf samples from co-expression of *CsupYPAQ, CsupKPSE* and *CsupFAR2* also contained an albeit very small but still significant amount of Z11-16:Ald (Fig. 7b). This implied the existence of an endogenous plant activity that works on Z11-16:OH for producing Z11-16:Ald.

**Figure 7.**
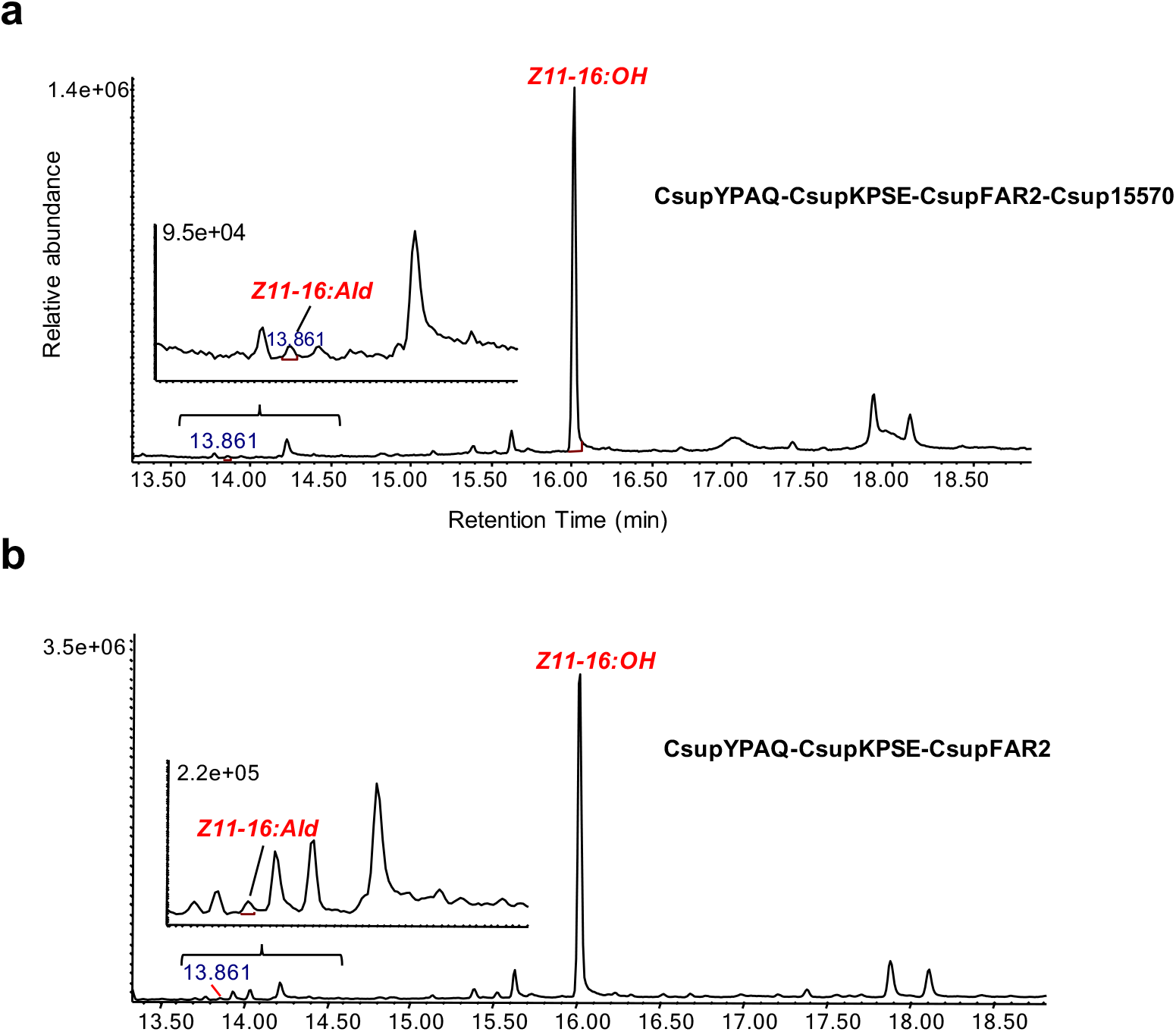
Heterologous expression of fatty alcohol oxidase candidate *Csup15570* with desaturase genes *CsupYPAQ* and *CsupKPSE* and reductase gene *CsupFAR2* from *Chilo suppressalis* in *Nicotiana benthamiana*. GC/MS analysis of Z11-16 fatty alcohol and aldehyde from plant leave expressing a) *CsupYPAQ*-*CsupKPSE-CsupFAR2*-Csup15570, b) *CsupYPAQ*-*CsupKPSE-CsupFAR2*. Fatty alcohol and aldehyde are shown in red.

### Transiently genetically modified *Nicotiana benthamiana* releasing moth pheromones

The newly identified desaturase and reductase genes with desired properties were used together with two genes previously identified, to produce plants transiently genetically modified to release moth pheromone compounds. *Nicotiana benthamiana* plants infiltrated with the functional Δ11 desaturases gene *AtrΔ11* and thioesterase gene *CpuFatB1*, and the newly identified desaturase and reductase genes *CsupYPAQ* and *CsupFAR2* (Fig. 8a), released a mixture of Z11-16:OH and Z11-16:Ald that could be collected as volatiles from the plant leaves (Fig. 8c-d). Upon co-expression of the acetyltransferase gene *ATF1*, Z11-16:OAc was also released together with Z11-16:OH and Z11-16:Ald. In addition, to explore the possibility of increasing the amount of moth pheromone compounds released from plant leaf, a *N. tabacum* trichome specific promoter *pCYP71D16* was used to control the expression of acetyltransferase gene *ATF1*. Consequently, the plant released significantly higher amounts of Z11-16:OAc, as well as Z11-16:Ald and Z11-16:OH, compared to when *ATF1* expression was controlled by the constitutive promoter *p35S* (Fig. 8d). The accumulation of Z11-16:Ald and Z11-16:OH were also higher in the leaves when *ATF1* was controlled by *pCYP71D16*, while Z11-16:OAc accumulation in the leaves showed no significant difference between the two strategies (Fig. 8e).

**Figure 8.**
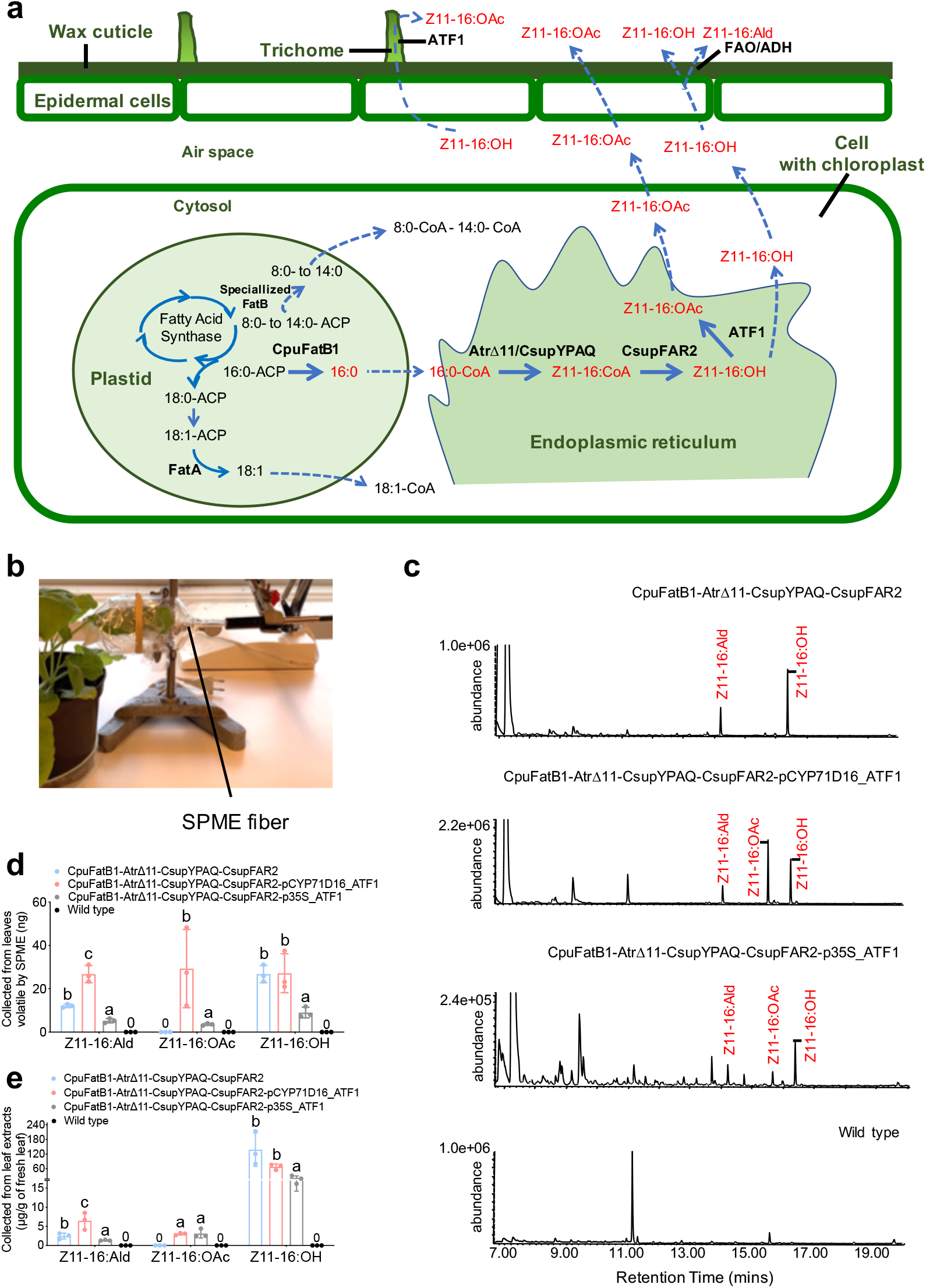
Rapid assembly of pheromone biosynthetic pathway in *Nicotiana benthamiana* for the production and release of moth sex pheromone components. The Cauliflower mosaic virus 35S promoter (*p35S*) and Octopine Synthase gene terminator (*tOCS*) have been used to regulate gene expression in plants. *ATF1* has also been controlled by trichome specific promoter *pCYP71D16*. (a) Step-wise metabolic engineering strategy for leaf-based pheromone production of (*Z*)-11-hexadecenol, (*Z*)-11-hexadecenal and (*Z*)-11-hexadecenyl acetate. (b) Solid-phase microextraction (SPME) of headspace volatiles from a genetically modified *N. benthamiana* leaf four days after infiltration. (c) GC-MS chromatograms (INNOWax column) of volatiles collected during 48 hours from an infiltrated *N. benthamiana* leaf. (d) The amount of collected pheromone compounds from released volatiles. (e) The amount of pheromone compounds from leaf extracts. Unpaired *t*-test and two-way ANOVA were used for the statistical analysis. P < 0.05 was considered as statistically significant. The error bars represent the standard errors of the mean (S.E.M.).

## Discussion

Enabling plants to release heterologously produced moth pheromones is an essential prerequisite for developing an IPM strategy in which the “pheromone-releasing” plants can be used for mating disruption of pests or to selectively attract a specific pest insect. We successfully made *N. benthamiana* release the major pheromone compound, Z11-16:Ald, of *C. suppressalis* and the common pheromone compounds Z11-16:OH and Z11-16:OAc, by transient expression of the functionally characterized *C. suppressalis* Δ11 desaturase *CsupYPAQ*, the fatty acyl reductase *CsupFAR2*, and the yeast acetyltransferase gene *ATF1*. Furthermore, characterization of the highly active pheromone biosynthetic genes from *C. suppressalis* is an important addition to the genetic toolbox for constructing heterologous platforms to produce customized insect pheromones.

Understanding the mechanisms underpinning moth pheromone biosynthesis is considered the basis for developing biological factories for moth pheromone production. In the present study, we demonstrate the key roles of three genes in the biosynthesis of the *C. suppressalis* sex pheromone, i.e. a Δ11 desaturase *CsupYPAQ* that acts specifically on palmitic acid to produce Z11-16:acid; a Δ9 desaturase *CsupKPSE* producing Z9-16:acid, the precursor of the minor pheromone component from palmitic acid; and *CsupFAR2* that transforms the acid precursors into the corresponding fatty alcohols. The enzyme(s) transforming the alcohols into the final aldehyde pheromone components remains elusive (Fig. 9).

**Figure 9.**
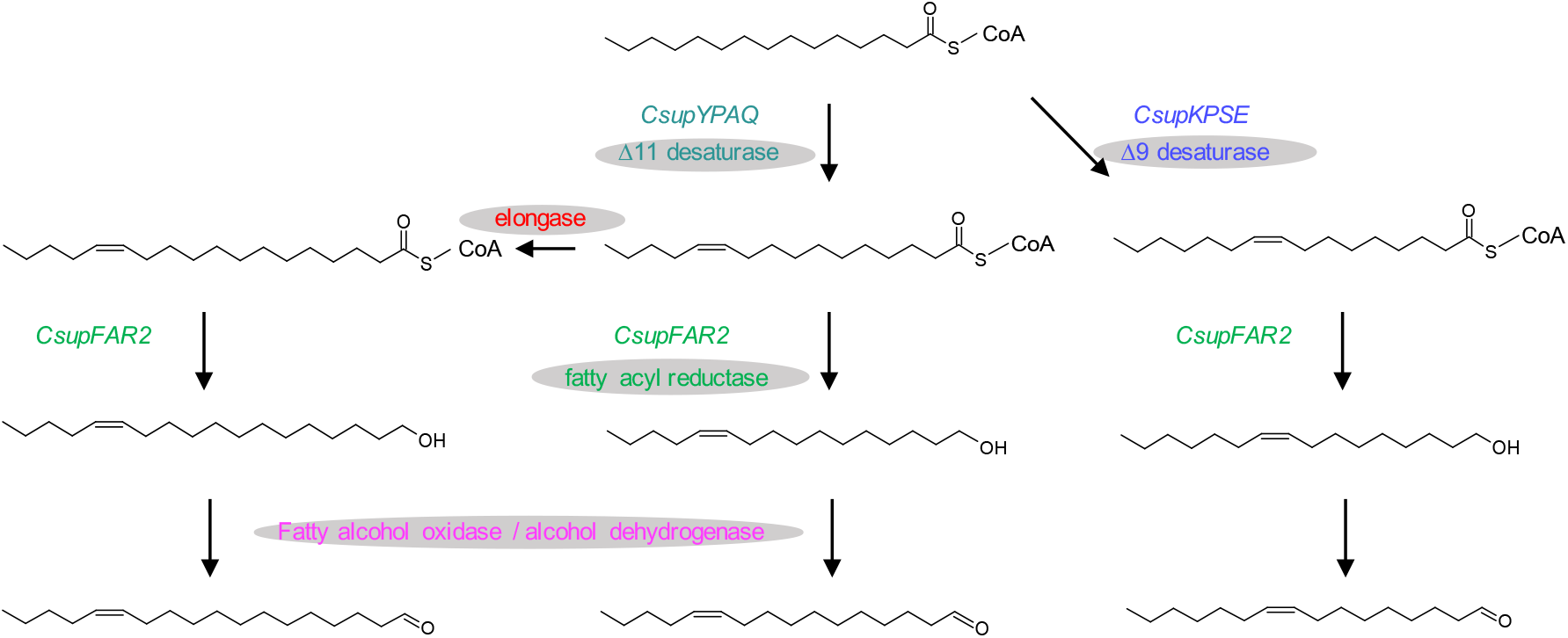
Biosynthetic pathways towards the sex pheromone of *Chilo suppressalis* and the key genes involved in each step.

In insects, the acyl-CoA desaturase gene family has had a very dynamic evolutionary history (Roelofs et al. 2003; Helmkampf et al. 2015; Knipple et al. 2002). As a result of gene duplication bursts, many opportunities have arisen for evolution to explore protein space and produce proteins with unique and novel functions. In phylogenetic analysis, *CsupYPAQ* clusters in the Lepidoptera-specific XXXQ/E clade (Fig. 1), which is a lineage that is composed of pheromone biosynthetic desaturases with diverse specificities (Knipple et al. 2002). With the exemption of the Δ9 desaturase from the tortricid species *Cydia pomonella* (Lassance et al. 2021), the FADs clustering in this clade are Δ11/Δ10 FADs but in some cases they have additional functions and are then classified as multifunctional. For example, *Trichoplusia ni* and *Spodoptera exigua* from Noctuoidae using Δ11 (Knipple et al. 1998), and Δ11/Δ12 (Xia et al. 2019) desaturations, respectively; *Thaumetopoea pityocampa* from Notodontidae using Δ11/Δ13 desaturations (Serra et al. 2007); *Bombyx mori* from Bombycidae using Δ11/10,12 desaturations (Moto et al. 2004); *Epiphyas postvittana, Choristoneura rosaceana, Agryrotaenia velutinana* from Tortricidae using Δ11 desaturations (Liu et al. 2002a; Liu et al, 2002b; Hao et al. 2002b), from the same family *Planotortrix octo* using Δ10 desaturation (Hao et al. 2002a) (Fig. 1). Interestingly, even though *CsupYPAQ* clusters close to *OfuZ/E11* (*Ostrinia furnacalis*) and *OnuZ/E11* (*O. nubilalis*) from Crambidae species, which produce both *Z* and *E* isomers of Δ11-tetradecenoic acid (Roelofs et al. 2002), *CsupYPAQ* specifically produces (*Z*)-11-hexadecenoic acid. In addition, *CsupYPAQ* shares 59.7% and 61.8% of amino acid identity to *SexiDes5* (*S. exigua*) and *SlitDes5* (*S. litura*), respectively, which are also both Δ11 FADs and have a very wide substrate preference, from 14 to 18 carbon chain length fatty acids (Xia et al. 2019). All evidence at hand advocates for caution when predicting gene functions based on homology to currently characterized gene sequences, and functional testing is essential in terms of enhancing our mechanistic understanding of the relation between sequence and function of desaturases. Ding et al. (2016) reported that one amino acid (258E) at the cytosolic carboxyl terminus of the protein is critical for the *Z* activity of the *C. rosaceana* FAD. The relative specificity of *CsupYPAQ* makes it an interesting candidate to further explore the correlation between the sequence and function of FADs by using site-directed mutagenesis and functional testing. Functional assays of *CsupYPAQ* showed similar results in the yeast and plant platforms but *CsupKPSE* expression did not yield oleic acid in *N. benthamiana* while it did in *S. cerevisiae*. Phylogenetically, *CsupKPSE* clustered in the ‘Δ9, C_16_>C_18_’ clade, however, it showed C_18_>C_16_ function when oleic acid was absent in the *ole1*/*elo1* yeast.

Fatty acyl reductases are used for converting fatty acyls into their corresponding alcohols during moth pheromone biosynthesis. Sometimes these FARs are referred to with the epithet “pheromone-gland specific” (pgFAR) although specific expression in the pheromone gland is not evident. Phylogenetically, the pgFAR clade containing the genes seems to be specific to Lepidoptera (Löfstedt et al. 2016). To date, several genes encoding pgFARs have been characterized in moth species such as *Bombyx mori* (Moto et al. 2003), *O. nubilalis* (Lassance et al. 2010), *Heliothis armigera* (Hagström et al. 2012), and *Spodoptera* spp (Antony et al. 2016). In this study, *CsupFAR2* is confirmed to be involved in sex pheromone biosynthesis of *C. suppressalis*. Previous studies have demonstrated the interplay between the abundance ofthe pheromone fatty acyl precursors and pgFARs in shaping the pheromone composition (Hagström et al. 2012). Although *CsupFAR2* shows activity on C_16_-C_18_ fatty acid substrates, it is interesting that *CsupFAR2* shows high selectivity for the Z11-16:acid when both Z11-16:acid and Z9-16:acid are present as substrates (Fig 6d and 6i). Taking into account the expression of *CsupYPAQ* and *CsupKPSE* in combination with *CsupFAR2*, the substrates preference of *CsupFAR2* suggesting that the FADs and FARs are both engaged and of importance in shaping the ratio of pheromone components in *C. suppressalis*.

Fatty alcohols may be pheromone components for many moth species but often fatty alcohols will be transformed to aldehyde pheromone components (Teal and Tumlinson 1986; Fang et al. 1995; Hoskovec et al. 2002). Since the 1980s, studies about moth aldehyde pheromone biosynthesis are diverging and non-conclusive (Morse and Meighen 1984; 1986; Teal and Tumlinson 1986; Foster et al. 2019; Jurenka 2021). The enzyme that produces pheromone aldehydes, and thus the gene encoding it, has not been characterized. We heterologously expressed six fatty alcohol oxidases (FAO) /dehydrogenases (ADH)-like genes (Table 1) in *N. benthamiana* leaves but did not get any conclusively results indicating oxidation by any of the candidates. We observed a small amount of Z11-16:Ald in the leaf extracts when Z11-16:OH was produced in large amounts but this occurred also without heterologous expression of FAO/ADH candidates (Fig. 7 and Table 2). The aldehyde production, in this case, may thus be due to an endogenous FAO/ADH activity from *N. benthamiana*. At the same time, it is interesting to note that all infiltrations of *N. benthamiana* leaves with functional FAD and FAR genes, with or without the acetyltransferase, made the plant release a remarkable amount of Z11-16:Ald as volatiles (Fig. 8c-d). The same infiltrated leaves contained very small amounts of Z11-16:Ald in the leaf extracts (Fig. 8e). This result provides evidence from a plant to support the hypothesis of Foster and Anderson (2019) that in moth females the alcohols and aldehydes are produced and/or stored in different compartments of the gland. In *H. virescens*, they found the aldehydes mainly on the cuticle of the gland whereas the alcohols were found inside the gland beneath the cuticle. In our study, Z11-16:Ald was abundant in the volatiles and less so in the leaf extracts, which implies that the alcohol component Z11-16:OH produced by pgFAR might have been converted to aldehyde when or after it went through the leaf wax cuticle (Fig. 8a).

Acetate pheromone compounds are postulated to be produced by an acetyltransferase from its alcohol precursor. Until now, the genes encoding moth acetyltransferases involved in pheromone biosynthesis have escaped identification. Previously, a plant-derived gene *EaDAcT* that produces 3-acetyl-1,2-diacyl-sn-glycerols (acetyl-TAG) by the acetylation of diacylglycerol and acetyl-CoA (Durrett et al. 2010) was heterologously expressed in *N. benthamiana* for the production of C_14_-C_16_ acetate pheromone compounds with significant but poor activity (Ding et al. 2014). Recently, Mateos-Fernández et al. (2021) also expressed *EaDAcT* with *AtrΔ11* and *HarFAR* in *N. benthamiana*, which resulted in the production and release of Z11-16:OH and Z11-16:OAc. In the present study, the *ATF1* which is naturally responsible for the synthesis of volatile esters that contribute to the fruity aroma of fermented alcoholic beverages (Verstrepen et al. 2003) from *S. cerevisiae* was co-expressed with moth FADs and a FAR in *N. benthamiana*. It resulted in the production of a high amount of acetate (Fig. 8e) compared to the previous study with *EaDAcT* (Ding et al. 2014). This result is consistent with activity differences in production of acetate pheromone compounds when *ATF1* and *EaDAcT* were expressed in *S. cerevisiae* (Ding et al. 2016).

Efficient transient expression of insect genes in plant leaves for pheromone production was demonstrated by Ding et al. (2014). Plant-produced pheromones or immediate precursors have been confirmed to perform favourably compared to conventionally chemistry-produced pheromones for trapping male moths (Ding et al. 2014; Xia et al. 2021, accepted) and are ready to be introduced for pest management (Löfstedt and Xia 2021). “Pheromone plants” engineered for the production of insect pheromones may be applied for pest control following at least two different strategies. One strategy is the production of pheromones/pheromone precursors in the plants, isolation and down-stream processing of the target compounds when the plant tissue has been harvested, for subsequent use of the active ingredient for moth trapping and mating disruption. The other strategy involves developing genetically modified plants capable of releasing pheromone compounds into the environment and, the pheromone-producing plant serves as a dispenser and may directly influence the behaviour of insects in the environment.

Optimization of plants as producers and actual dispensers of insect pheromones require different measures compared to engineering of plants for efficient storage of the same compounds or their precursors. For acetate release, not only a functional acetyltransferase has to be present to produce acetate, but the product has to be released, which makes it important to understand the secretion mechanism of plant volatiles. In a large number of plants, trichomes that are tiny specialized hair structures for secondary metabolite production, play a prominent role for release of volatiles. Biosynthesis of plant diterpenes occurs in trichome heads, where secretory vesicles and cells are located (Kandra and Wagner 1988; Duke 1994; Guo et al. 1995). We found that the plant carrying *ATF1* controlled by *pCYP71D16*, a trichome-specific promoter cloned from *N. tabacum*, released 3 to 10 fold higher amounts of Z11-16:OAc and also much higher amounts of Z11-16:Ald and Z11-16:OH compared to when expressed from *p35S* (Fig. 8d). This suggests that release of pheromone compounds benefits from expression of the pheromone biosynthetic genes in the trichomes. More pheromones might be released along with the native plant volatiles when they are produced in trichomes.

Following several proof-of-concept studies demonstrating the possibility of producing moth pheromones in plants and in microbial cell factories, we showed that plants can be engineered to actually release the pheromone compounds, paving the way for stable transformation of plants to be used in different IPM strategies.

## Materials and Methods

### Sequences and phylogenetic analysis

The open reading frames (ORFs) of the FAD-like genes *CsupYPAQ* (GenBank accession number: MN453822) and *CsupKPSE* (MN453823), the FAR-like gene *CsupFAR2* (MN453825) and the FAO/ADH-like genes *CsupFAO_15570, CsupFAO_9572, CsupADH_10975, CsupADH_14583* and *CsupADH_17286* were obtained from the *C. suppressalis* pheromone gland transcriptome sequences (Xia et al. 2015). The ORF of the ADH-like gene *HzeaADH7* was obtained from the transcriptome sequences reported in Dou et al. 2019. *CsupYPAQ, CsupKPSE, CsupFAR2* correspond to *CsupDes4, CsupDes1*, and *CsupFAR2* in Xia et al. (2015). The *AtrΔ11* (JX964774) and *CpuFatB1* (AGG79283) were amplified from entry clones (Ding et al. 2014). The promoter *pCYP71D16* was cloned from *Nicotiana tabacum* genome as described in Wang et al. (2002).

The phylogenies of the FAD and FAR sequences were constructed using *C. suppressalis* FADs and FARs with the functionally characterized sequences retrieved from the GenBank from National Center for Biotechnology Information (NCBI) (https://www.ncbi.nlm.nih.gov/). The Neighbor-Joining tree was constructed using MEGA version 5.0 (Tamura et al. 2011). The bootstrap consensus tree inferred from 500 replicates was taken to represent the evolutionary history of the analysis of the genes.

### Cloning of gene candidates for functional assay

All the gene candidates were synthesized by Invitrogen, Life Technologies. PCR amplification of each gene candidate was performed using the synthesized sequence as the template with a pair of specific primers (Table S1) with attB1 and attB2 sites incorporated on a Veriti Thermo Cycler using Phusion Flash High-Fidelity PCR Master Mix (Thermo Scientific ™). Cycling parameters were as follows: an initial denaturing step at 98 °C for 30 s, 38 cycles at 98 °C for 5 s, 55 °C for 10 s, 72 °C for 50 s, followed by a final extension step at 72 °C for 10 min. The PCR products were subjected to agarose gel electrophoresis and purified using the GeneJET Gel Extraction Kit (Thermo Scientific™). Then the ORFs were cloned into the pDONR221 vector in presence of BP clonase (Life Technologies) to generate the entry clone. After the entry clone for each ORF was confirmed by sequencing with M13+ and M13-primers, it was cloned into either the yeast expression vectors pYEX-CHT or pYES52 or the plant expression vector pXZY393 by LR reaction (Invitrogen). The resulting expression clones were analyzed by sequencing.

### Yeast heterologous expression

The experimental workflow is shown in Fig. 10a. The expression clones containing *CsupYPAQ* and *CsupKPSE* were introduced into the double deficient *ole1/elo1* strain (*MATa elo1::HIS3 ole1::LEU2 ade2 his3 leu2 ura3*) of the yeast *S*.*c*., while the expression clones containing *CsupFAR2, CsupYPAQ*-*CsupFAR2*, pYEX-CHT empty vector and pYES52 empty vector were introduced into the *INVSc* strain (*MATa HIS3 LEU2 trp1-289 ura3-52*), using the *S*.*c*. easy yeast transformation kit (Life technologies). For selection of uracil prototrophs, the transformed yeast was allowed to grow on SC plate containing 0.7% of YNB (w/o aa, with ammonium sulfate) and a complete drop-out medium lacking uracil (Formedium™ LTD, Norwich, England), 2% of glucose, 1% of tergitol (type Nonidet NP-40, SigmaeAldrich Sweden AB, Stockholm, Sweden), 0.01% of adenine (Sigma), and containing 0.5 mM Z10-17:Me (Sigma) as extra fatty acid source (no supplementary for *INVSc* strain).

**Figure 10.**
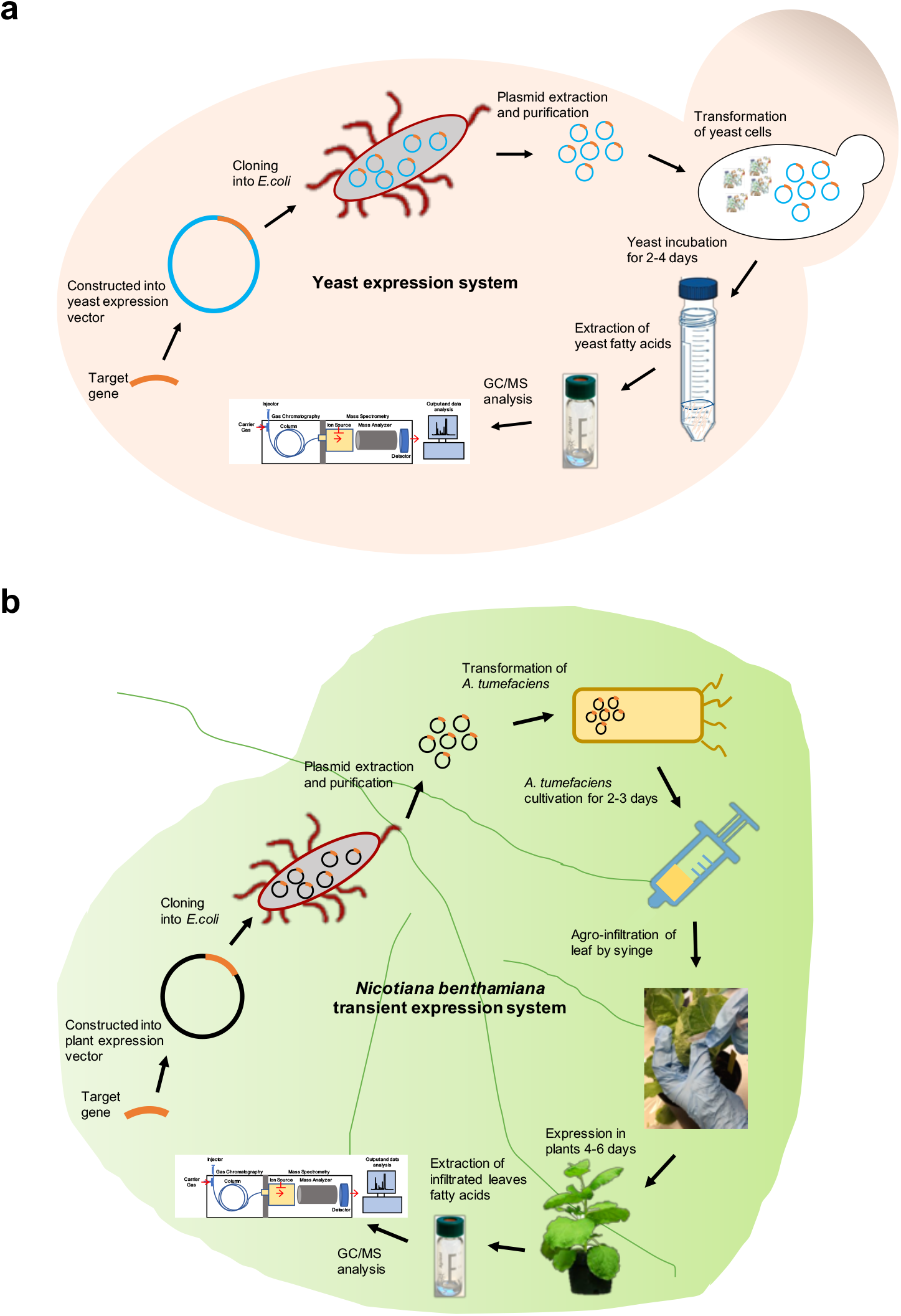
Experimental workflow of the heterologous expression in a) yeast, and b) plant. Adapted from Löfstedt and Xia (2021).

After 4 days (2 days for *INVSc* strain) incubation at 30 °C, individual colonies were picked to inoculate 2 mL selective medium and then grown at 30 °C and 300 r.p.m for 48 h. Yeast cultures were diluted to an OD600 of 0.4 in 5 mL fresh selective medium containing 0.5 mM CuSO_4_ (for pYEX-CHT vector) or 10% galactose (for pYES52 vector) for induction. Then the yeast cells (medium was also harvested for *CsupFAR2* and *CsupYPAQ*-*CsupFAR2* yeast line) were harvested after 48 h incubation in a shaking incubator at 30 °C for fatty acid or alcohol analysis.

### *Nicotiana benthamiana* material and growth condition

The wild type *N. benthamiana* plants for the *Agrobacterium* infiltration were grown in the greenhouse under 16 h/8 h light conditions. The greenhouse’s growth temperature and relative humidity were set at 24 °C/18 °C day/night and 40% RH.

### *Agrobacterium* infiltration of *Nicotiana benthamiana*

The experimental workflow is shown in Fig. 10b. The genes transformed for pheromone production in plants were generally controlled by the Cauliflower mosaic virus 35S promoter (*p35S*) and Octopine Synthase gene terminator (*tOCS*) but *ATF1* was in one version also controlled by the tobacco trichomes specific promoter *pCYP71D16*. The expression clones containing *CsupYPAQ, CsupKPSE, AtrΔ11, CpuFatB1, ATF1, CsupFAR2, CsupFAO_15570, CsupFAO_9572, CsupADH_10975, CsupADH_14583, CsupADH_17286* and *HzeaADH7* were introduced into *Agrobacterium tumefaciens* GV3101 strain (MP90RK) by electroporation (1700 V mm^-1^, 5 ms, Eppendorf 2510). A viral silencing suppressor protein *P19* was introduced into GV3101 strain as well in order to inhibit the host cells’ transgene silencing apparatus and extend transgene expression over a longer period of time with a higher degree of expression (Canto et al. 2006).

The *Agrobacterium* infiltration of *N. benthamiana* started with cultivation of 10 mL of *Agrobacterium* that contained individual gene constructs at 30 °C with LB medium supplemented with appropriate antibiotics overnight in a 300 r.p.m. incubator. Then 100 µM acetosyringone was added to the culture and grown for an additional 2-3 h to induce virulence genes encoded by *Agrobacterium* genome. Subsequently, the *Agrobacterium* were spun down at 4200 g for 5 min at room temperature and resuspended in infiltration buffer (5 mM MgCl_2_, 5 mM 4-morpholineethanesulfonic acid, 100 µM acetosyringone, pH 5.7). Then the optical density under 600 nm wavelength (OD_600nm_) of each *Agrobacterium* culture was measured to adjust final concentration of each culture to OD_600nm_=0.2, in a total volume of 20 mL infiltration buffer as described before.

Afterwards, each final mixture of *Agrobacterium* cells was drawn up into a 1 mL syringe without needle and infiltrated into the underside of a suitable four-week-old *N. benthamiana* leaf, with a gentle squeeze on the plunger and modest pressure on the leaf using a finger. By this the *Agrobacterium* solution was forced into the mesophyll spaces wetting the leaf. Five entire leaves of similar age from randomly selected four-week-old plants were infiltrated. Then, plants were maintained in the growth chamber for four days with sufficient watering.

### Lipid extraction and preparation

For yeast lipid analysis, all the cells and media (for alcohol analysis) were extracted by 1 mL of methanol/chloroform (2:1, v/v) and then 1 mL of water was added to produce a biphasic mixture, which was then vortexed vigorously and centrifuged at 2000 r.p.m for 2 min. Then ca. 330 µL of the chloroform phase containing the total lipids and pheromone compounds was transferred to a new glass vial, followed by evaporation to dryness under a gentle flow of nitrogen. The residues were used for fatty acid and pheromone analysis. For fatty acid analysis, 1 mL of 2% sulphuric acid in methanol was added, and the sample was incubated 1 h at 90°C for methanolysis to occur. Subsequently, 1 mL water and 1 mL heptane were added and the mixture was vortexed energetically. Finally, ca. 1 mL of heptane phase containing the fatty acids in the form of corresponding methyl esters was transferred to a new glass vial for GC/MS analysis. For pheromone compounds analysis, 200 µL heptane was added after the evaporation to dryness and after vortex vigorously the sample was then transferred to a new GC/MS analytical glass vial.

For leaf lipid analysis, ca. 100 mg of fresh leaf tissue from each sample was collected, and 3.12 µg internal standard methyl nonadecanoate (19:Me) or 1.8 µg internal standard (*Z*)-8-tridecenyl acetate (Z8-13:OAc) was added per gram fresh leaf to the samples for fatty acid and pheromone compound analysis respectively, following the same protocol as described above.

### Sampling volatile compounds in static plant headspace

The set up for plant static headspace volatile collection is shown in Fig. 8b. After 3-5 days of infiltration, the plant was used for the collection of volatiles. The infiltrated plant leaf was enclosed in a glass funnel and covered by a transparent and odorless oven bag, and a solid phase microextraction (SPME) fiber (65 µm film thickness, polydimethylsiloxane/divinylbenzene (PDMS/DVB), Supelco, Bellefonte, PA) was inserted from the stem of the funnel. The volatiles were collected for 24 h before GC/MS analysis. The funnel was washed with ethanol and acetone between collections. The collections for each treatment had at least three biological replicates. Synthetic Z11-16:OH was used as external standard to quantify the target compounds.

### Gas chromatography/mass spectrometry (GC/MS)

Yeast and plant-leaf samples were analyzed using an Agilent 5975 mass selective detector coupled to an Agilent 6890 series gas chromatograph equipped with a polar column (HP-INNOWax, 30m x 0.25 mm, 0.25 µm film thickness) or using an Agilent 5975C mass selective detector coupled to an Agilent 7890A series gas chromatograph equipped with a non-polar column (HP-5MS, 30m x 0.25 mm, 0.25 µm film thickness). The compounds were identified based on mass spectra and retention times on two columns being identical to synthetic standards. Helium was used as carrier gas (average velocity 33 cm/s). The injector was configured in splitless mode at 250 °C. The oven temperature was set at 80 °C for 1 min, then increased to 230 °C at a rate of 10°C/min and held for 10 min.

DMDS derivatization was performed to determine the position of double bonds in target compounds, according to Dunkelblum et al. (1985). The DMDS-adducts were analyzed by GC/MS equipped with the non-polar column (HP-5MS) under the following oven temperature program: 80 °C for 2 min, then increased at a rate of 15 °C/min to 140 °C, and then increased at a rate of 5°C /min to 260°C, and held for 3 min.

### Chemicals

Fatty acids references and synthetic pheromone components of various origins were available from our laboratory collection of pheromone compounds.

### Statistical analysis

The data collected for statistical analysis are from three independent experiments. Each experiment has four to five biological replicates. Data were processed by Prism version 8.0. Unpaired *t*-test and two-way ANOVA were used for the statistical analysis. P < 0.05 was considered as statistically significant.

## ACKNOWLEDGEMENT

This work was supported by funding from the Swedish Foundation for Strategic Research (No. RBP 14-0037, Oil Crops for the Future), the European Union’s Horizon 2020 research and innovation programme (No. 760798, OLEFINE), Formas (No. 2010-857 and 2015-1336), and the Carl Trygger Foundation for Scientific Research (No. CTS 14:307 and CTS KF17:15) to CL, and the Jörgen Lindström’s Scholarship Fund and the Royal Physiographic Society in Lund to YHX. The Chinese Scholarship Council supported Yi-Han Xia’s PhD scholarship. We thank Carin Jarl-Sunesson for helpful advice on plant cultivations. We thank Erling Jirle for excellent technical support. PH recognizes support from the strategic research program Trees and Crops for the Future (TC4F).

## Author contribution statement

YHX, BJD, and CL conceived the study. YHX and BJD carried out vector design and sequencing. YHX performed gene functional assays. YHX performed plant cultivation and sample analysis. HLW contributed to plant volatile collections. BJD, HLW, PH, and CL provided technical guidance and suggestions on metabolic engineering strategies. YHX drafted the manuscript with assistance from CL, all authors edited the manuscript and approved the final version.

