## Supplemental Table 1 for "Making Plants Smell Like Moths: *Nicotiana benthamiana* release moth pheromone alcohol, aldehyde and acetate upon transient expression of biosynthetic genes of different origin"

**Table S1. Primers used in this study.**

| Gene abbreviation | Primer sequence (5’-3’) |
|---|---|
| *CsupYPAQ_F* | GGGACAAGTTTGTACAAAAAAGCAGGCTTAATGGCCCCGAATTCAATTCAAAATG |
| *CsupYPAQ_TT_79R* | CTAACATAACTATAAAAAAATAAATAGGGACCTAGACTTCAGGTTGTCTAACTCCTTCCTTTTCGGTTAGAGCGGATTCAATCCTCCGTCTTGCTGTAC |
| *TT_145R* | GTTACATGCGTACACGCGTCTGTACAGAAAAAAAAGAAAAATTTGAAATATAAATAACGTTCTTAATACTAACATAACTATAAAAAAATAAATAG |
| *TT_attB5r_R* | GGGGACAACTTTTGTATACAAAGTTGACTCTTCGAGCGTCCCAAAACCTTCTCAAGCAAGGTTTTCAGTATAATGTTACATGCGTACACGCGTCTG |
| *attB5_Gal1_ F* | GGGGACAACTTTGTATACAAAAGTTGTAACGGATTAGAAGCCGCCGAGCGGGTGAC |
| *Gal1_CsupFAR2_F* | ACGTCAAGGAGAAAAAACCGTGGTTGTGAAAATGGAAC |
| *Gal1_CsupFAR2_R* | GTTCCATTTTCACAACCACGGTTTTTTCTCCTTGACGT |
| *CsupFAR2_404F* | CTCTGTCCGAGGCTATCATCATC |
| *CsupFAR2_1257R* | CTTGTCGATGAACAAGAACTGGTTG |
| *Gal1_94F* | GATGTGCCTCGCGCCGCACTGC |
| *Gal1_395R* | GAGGTATATTAACAATTTTTTGTTGATAC |
| *CsupELO1_F* | GGGGACAAGTTTGTACAAAAAAGCAGGCTAATGGAGGTGCTAAGGAGACTAG |
| *CsupELO1_R* | GGGGACCACTTTGTACAAGAAAGCTGGGTTTTACTGGGACGCCACCGCTCCTGCCATC |
| *CsupELO3_F* | GGGGACAAGTTTGTACAAAAAAGCAGGCTAATGAACGGTGCTAATACGACCTTCGAAATGTC |
| *CsupELO3_R* | GGGGACCACTTTGTACAAGAAAGCTGGGTTCTAGCAATCTTTTGCTTTTCCATTGGCTG |
| *CsupELO4_88F* | CTGATCATCTGCCTGTCCTACGTG |
| *CsupELO4_783R* | CTTAGCGCGCACTTTGGTCTTGG |
| *Csup15570_605F* | GATCGAAAATTTTCAAATTCCTTCATTC |
| *Csup15570_1550R* | CAAGTCTTCCTCATATTCAAAGTAG |
| *Csup9572_553F* | AGGAATTCGACTTATCACGG |
| *Csup9572_948R* | CATCAACAATTGGGCAGATC |
| *Csup10975_295F* | GCTAAAGTGAAGTCCCTGAA |
| *Csup10975_625R* | TGATCAGGATCTTGTTAGCG |
| *Csup14583_170F* | GTCACACTGACGCATATACT |
| *Csup14583_633R* | ACCAAAGATTGCACAATTGG |
| *Csup17286_307F* | GTTGCTACAGGTTTACATGC |
| *Csup17286_1097R* | TATAGTGCCTCAATGCCATC |
| *HzeaADH7_37F* | GAACCACACCAAGAAGATCT |
| *HzeaADH7_R* | ATCACGTAGTTCTCGTTAGC |
